# Viral gain-of-function experiments uncover residues under diversifying selection in nature

**DOI:** 10.1101/242495

**Authors:** Rohan Maddamsetti, Daniel T. Johnson, Stephanie J. Spielman, Katherine L. Petrie, Debora S. Marks, Justin R. Meyer

## Abstract

Viral gain-of-function mutations are commonly observed in the laboratory; however, it is unknown whether those mutations also evolve in nature. We identify two key residues in the host recognition protein of bacteriophage λ that are necessary to exploit a new receptor; both residues repeatedly evolved among homologs from environmental samples. Our results provide evidence for widespread host-shift evolution in nature and a proof of concept for integrating experiments with genomic epidemiology.

---

Many viruses can expand their host range with a few mutations^1-3^ that enable the exploitation of new cellular receptors^2,3^. Such mutations may be the first steps toward an epidemic outbreak; this observation has driven an expansion of theoretical^4^, experimental and surveillance studies of host-range shifts in emergent pathogens, including avian influenza^5-8^, coronaviruses^9,10^, HIV^11^ and ebolavirus^12^. Ideally, laboratory evolution experiments could be used to accelerate our understanding of how viruses expand their host range. However, it is not clear whether viral evolution observed under the chemical and physical environments of a laboratory can faithfully inform us about viral evolution in nature, for at least two reasons. First, evolutionary trajectories might be sensitive to differences in environmental conditions between the laboratory and nature. Second, the number of evolutionary paths sampled in laboratory experiments is very small compared to natural virus diversity due to the enormous size of viral populations. Indeed, some researchers have called for the suspension of virus gain-of-function laboratory experiments^13^ on the grounds that they would tell us little about real-world viral evolution at the risk of constructing viral strains that are pandemic to new hosts. Here, we use a harmless virus, bacteriophage λ, to demonstrate how gain-of-function experiments can identify mutations that mimic those that occur in nature: we find that two amino acid residues that are critical for gain of function in the laboratory recurrently evolve in nature.

Typical laboratory strains of λ infect *Escherichia coli* by binding to the outer membrane protein LamB^14^, but the phage rapidly evolves in the laboratory to exploit a different membrane protein, OmpF^3^. Since OmpF is not the usual *E. coli* receptor for λ, these experiments are a proxy for the ability of the phage to switch hosts. The evolved gain-of-function phenotype in λ, OmpF^+^, involves multiple non-synonymous mutations in the host-recognition gene *J*. Each OmpF^+^ isolate had between 4 and 10 single nucleotide substitutions in *J*, and none had insertions or deletions (indels). 97% of the substitutions in 24 independently evolved OmpF^+^ λ phage occurred in the *J* protein^3^ between residues 960 —1132, which we call the ‘specificity region’ of *J* (Figure 1A).

**Figure 1.**
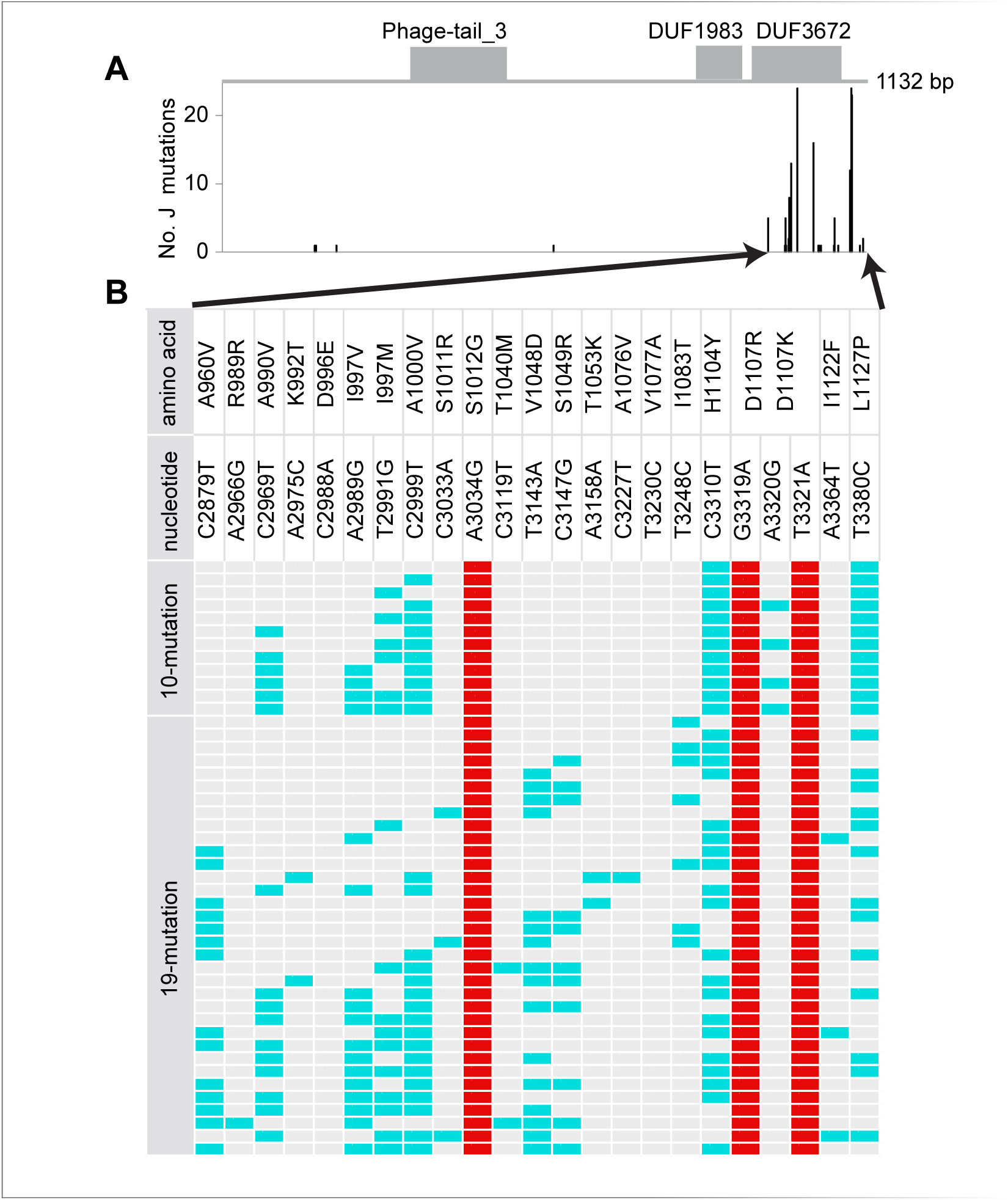
A) Distribution of J mutations evolved *en route* to OmpF^+^ and B) synthetic phage genotypes capable of using OmpF. A) Mutations summed across 24 genomes independently evolved to exploit OmpF in Meyer *et al*. (2012). Protein domains as annotated in the Pfam database are shown in gray. The majority of mutations either occur in the DUF3672 domain, or in the C-terminus past the annotated boundary of DUF3672. B) Synthetic OmpF^+^ genotypes indicated by colored cells (red marks the canonical three) observed after combinatorial engineering of 10 common mutations or 19 mutations when the canonical three were fixed. Amino acid and nucleotide changes are indicated by positions bookended by the wild-type state and then the evolved state. Amino acids in position 997 and 1107 have multiple derived states.

By comparing *J* among OmpF^+^ and OmpF^−^ λ, Meyer *et al*. (2012) suggested that the OmpF^+^ innovation required four mutations: one at residue 1012, two in the codon for residue 1107, and a fourth mutation somewhere between residues 990 to 1000^3^. In this study, we directly tested this hypothesis by using Multiplexed Automated Genome Engineering (MAGE)^15^ to construct two combinatorial genetic libraries of the *J* mutations that evolved in Meyer *et al*. (2012). We constructed the first library by focusing on 10 commonly evolved *J* mutations^3^, from which we sequenced 33 OmpF^+^ isolates. All strains possessed the target mutation at residue 1012 and both target mutations at residue 1107; by contrast, some strains lacked a fourth mutation between residues 990 and 1000 (Figure 1). We call the mutations at residues 1012 and 1107 the “canonical mutations”.

To test whether the three canonical mutations were sufficient to confer the OmpF^+^ phenotype, we constructed a synthetic phage with just these mutations. Even though the synthetic phage was viable, it proved unable to exploit cells without the ancestral LamB receptor, demonstrating that at least four *J* mutations were necessary for laboratory-evolved OmpF^+^ phage. To find what further mutations might be needed to confer the OmpF^+^ phenotype, we constructed a second MAGE library using the phage with the three canonical *J* mutations as the baseline, and random combinations of 19 other *J* mutations found in the OmpF^+^ λ evolved by Meyer *et al*. (2012). Screening this second library produced 88 OmpF^+^ isolates. One OmpF^+^ isolate had just a single extra mutation in amino acid position 1083 in addition to the three canonical mutations (Figure 1). However, the majority of the OmpF^+^ genotypes did not possess this specific mutation, signifying that its function could be substituted by other *J* mutations. In total, this experiment revealed that four mutations are sufficient to evolve the innovation, but only two specific amino acid changes (at residues 1012 and 1107) are universally required to access OmpF in the context of laboratory experiments.

We then asked whether natural J sequences reflected the host-shift evolutionary dynamics seen in our gain-of-function experiments. We collected and aligned full-length homologous J protein sequences from UniRef100^16^ (1,207 highly similar sequences). Most sequences were prophage uncovered in the genomes of their *Enterobacteriaceae* hosts, including bacterial genera *Escherichia, Salmonella, Citrobacter, Edwardsiella*, and even *Cronobacter*.

Recall that 97% of substitutions in OmpF^+^ gain-of-function experiments occur in the specificity region. Likewise, the natural J homologs had disproportionate variation here: 30% of the total amino acid variation occurred in the specificity region, despite only being 15% of the total length of J (Figure 2A). This nonrandom clustering of variation in the specificity region strongly suggests that J has experienced substantial diversifying selection on host attachment. Furthermore, the 17 residues engineered into the MAGE libraries were significantly more variable than randomly chosen groups of 17 residues from J (non-parametric bootstrap: *p* < 10^−5^) and from the specificity region 960-1132 (non-parametric bootstrap: *p* = 0.0054) showing the experiments had identified evolutionary hotspots. However, the specific 17 amino acid substitutions evolved in the laboratory were not common in natural homologs suggesting that the selection experiments had the resolution to predict where changes would evolve, but not the exact change (Supplementary Figure 1).

**Figure 2.**
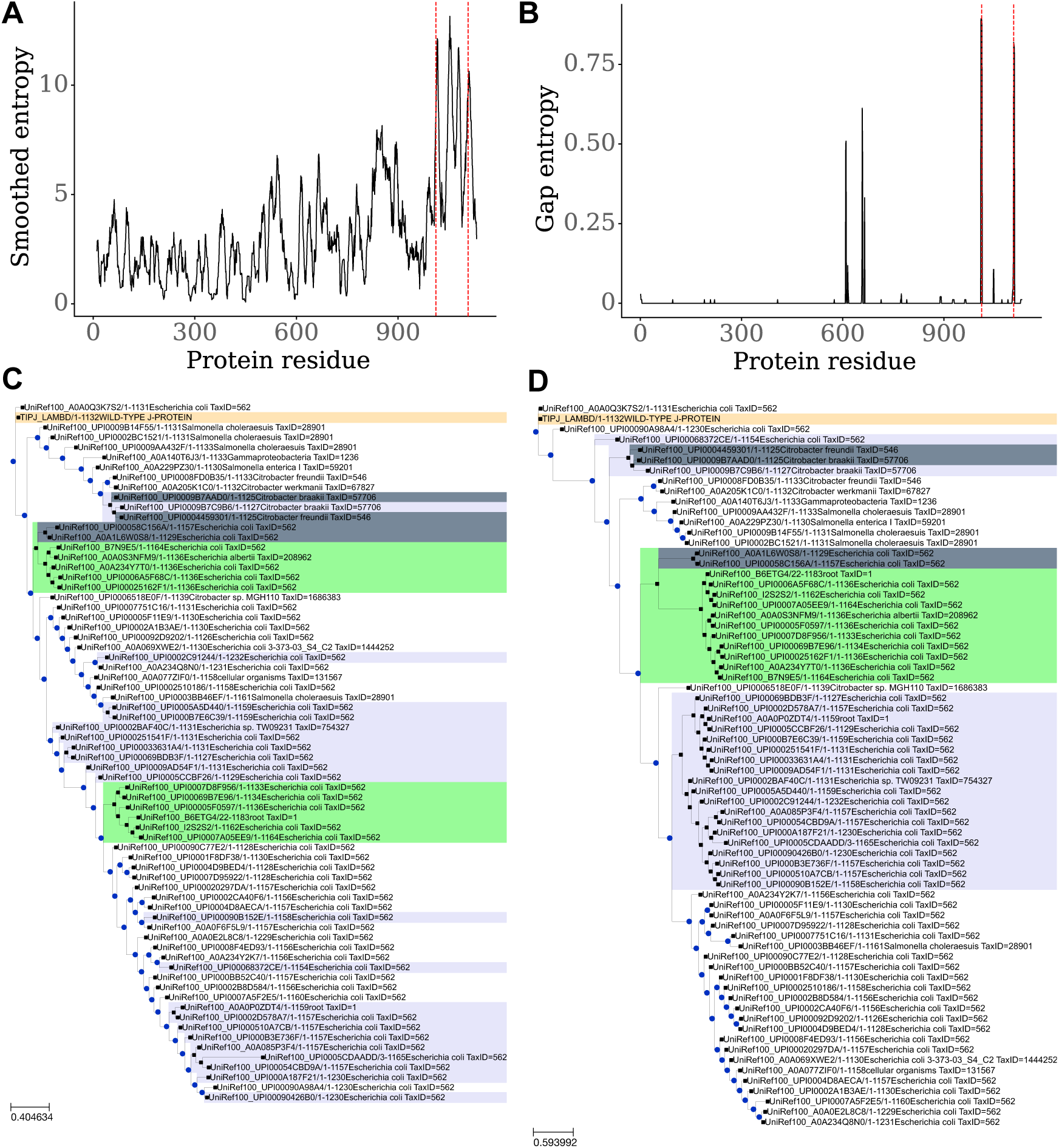
A) Sequence entropy averaged over a 10-residue window in J. Dashed red lines indicate residues 1012 and 1107. B) Gap entropy over the J protein. Dashed red lines indicate residues 1012 and 1107. C) Phylogeny for J protein. Only a subset of the operational taxonomic units (OTUs) are displayed: just OTUs with sequences ≥ 5% divergence from each other. The wild-type J protein sequence is labeled in tan. Clades with indels at residue 1012 are labeled in lavender. Clades with indels at residue 1107 are labeled in light green. Sequences with indels at both positions are labeled in light steel gray. D) Phylogeny for residues 960–1132 of J protein. OTUs have at least 5% divergence from each other, and are labeled as in (C).

Focusing in on the two residues critical for the OmpF^+^ gain of function, 1012 and 1107, we find that they alone are more variable than random pairs of sites in J (non-parametric bootstrap: *p* = 0.00101) and the specificity region (non-parametric bootstrap: *p* = 0.036). But what is more remarkable is the observation that in-frame indels are common in the regions around sites 1012 and 1107 (Figure 2B; Supplementary Data 1). We measured indel variation over J homologs using gap entropy: a two-state entropy counting each position in each aligned sequence as a gap or non-gap. Residues 1010–1013 and 1107–1109 have the highest gap entropy in J, ranging from 0.799 to 0.893 out of a maximum of 1.000 (Figure 2B; Supplementary Data 2). For comparison, the typical amino acid site in the J alignment had no indels (mode and median gap entropy = 0). That our experiments would correctly identify both regions with the most natural indel variation by chance is extremely unlikely (given the size of each region and the protein length 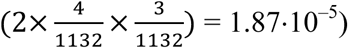. Indels can have large beneficial effects on proteins, including altering specificity by changing surface loops^14,17^, or causing structural rearrangements that improve function^18^.

To analyze J protein evolutionary dynamics in more depth, we built maximum-likelihood phylogenetic trees for J (Figure 2C) and its specificity region (Figure 2D). The trees are very different (normalized Robinson-Foulds distance = 0.94 out of 1.00), indicating that recombination has caused the history of the specificity region to differ from the history of the rest of the protein. This means that the specificity region is an evolutionary module that commonly recombines and circulates as a unit in the phage population. This observation supports the notion that this region has elevated rates of evolution and diversification.

Using the second phylogeny, we calculated site-specific evolutionary rates for the specificity region. The 17 residues studied in the MAGE experiments evolve faster than equally-sized random samples taken from the specificity region (non-parametric bootstrap, *p* = 0.0067) and residues 1012 and 1107 evolve faster than random pairs of sites sampled from the specificity region (non-parametric bootstrap, *p* = 0.0073).

Together, the accelerated rate of change and increased entropy at residues 1012 and 1107 suggest that these sites have experienced strong diversifying selection. Our combinatorial genetic studies showed that these particular residues determine receptor tropism, so it is reasonable to conclude that their unique evolution is due to host-range evolution. Mathematical theory has demonstrated that receptor-use evolution should lead to diversification since accessing new host types alters the phages’ ecological niches^19^. Indeed, diversification was observed in a related experiment where λ evolved to exploit two host cell types with different receptors^20^. To test whether indels at 1012 and 1107 cause diversification, we compared the branches where indels occur versus all others. If the indels cause ecological differentiation, then they should reduce intraspecific competition for hosts and facilitate the long-term maintenance of distinct evolutionary lineages. Indeed, the branches on which indels occur are significantly longer than the other branches of the specificity-region phylogeny (Kruskal-Wallis test, *p* < 10^−8^; visualized by the discrete indel clades colored in lavender and light green in Figure 2D).

The congruence we find between laboratory and natural evolution in λ contrasts with work showing that beneficial mutations in Richard Lenski’s long-term experiment anticorrelate^21^ with natural protein variation in *E. coli*. By design, Lenski’s experiment lacks the complex and variable selection pressures found in *E. coli’s* natural gut environment, leading the bacteria down evolutionary paths not taken in nature. By contrast, the dominant selection pressure in evolution experiments with bacteriophage— attachment to bacterial host cells— is probably also a dominant selection pressure on phage in the wild. Notably, another study on experimental evolution of bacteriophage has observed residues under positive selection in both the lab and nature^22^.

Together, these studies show that selection experiments on viruses cultured in the laboratory can inform evolution in nature. Based on the patterns of J sequence variation observed, we suggest that host-range evolution is common in this group of viruses, and perhaps others too. While the frequency of host-range evolution may be unsettling, our work also demonstrates potential methods to combat host shifts. In particular, worrisome mutations can be identified with functional genetic experiments as described here or with other laboratory techniques such as deep mutational scanning^23^. This information can be combined with genomic surveillance efforts underway^24,25^ to devise more effective disease management strategies to eradicate problematic strains.

## Methods

### MAGE *experiments*

We modified λ strain cI857 (provided by Ing-Nang Wang, SUNY Albany) integrated into the genome of HWEC106 (provided by Harris Wang, Columbia University). We engineered the J substitutions into λ by using MAGE^15,26^ on strain HWEC106. MAGE uses the λ-red recombineering system provided on the pKD46 plasmid^27^. For a description of the oligos used, see Supplementary Table S1.

For the first, 10-mutation library, we ran 18 rounds of MAGE and then screened for OmpF^+^ isolates. For the second library, we edited λ with oligos ‘a3034g’ and ‘g3319a t3321a’ to introduce the canonical three mutations. Next, we performed a number of different MAGE trials in order to maximize the number of unique alleles we observed. See Supplementary Table S2 for our MAGE strategy.

To screen the edited phage for OmpF^+^ genotypes, we induced the MAGE lysogen libraries and plated the phage on lawns of *lamB^−^ E. coli* strain JW3996 from the KEIO collection^28^. Plaques were picked from the lawns and the C-terminus of the J gene was Sanger sequenced at the Genewiz La Jolla, CA facility. Unpurified PCR products (Forward primer: 5’ CGCATCGTTCACCTCTCACT; Reverse primer: 5’ CCTGCGGGCGGTTTGTCATTT) were submitted.

### Analysis of natural *J* variation

We used the *evcouplings* pipeline^29^ to generate a jackhmmer^30^ alignment of 1,207 full-length J homologs (parameter settings: bitscore = 0.2; theta = 0.999; seqid_filter = 95 and 99 (thresholds for filtering highly similar sequences); minimum_sequence_coverage = 99 (require sequences to align to 99% of wild-type J protein); and minimum_sequence_coverage = 50. When analyzing the specificity region (say, constructing a phylogeny), we used residues 960-1132 of this full-length alignment.

We used RAxML^31^ to generate maximum-likelihood phylogenies using automatically selected amino acid substitution matrices (PROTGAMMAAUTO) based on model fit. To account for recombination when estimating site-specific evolutionary rates in the specificity region, we first ran a modified SBP algorithm^32^ on the specificity region. We found a putative recombination breakpoint at residue 1000, supported by Akaike’s information criterion but not the Bayesian information criterion. We therefore partitioned the specificity region at residue 1000 and re-calculated trees for each partition using FastTree^33^ with an LG substitution matrix^34^. We next calculated site-specific evolutionary rates with LEISR^35^, a scalable implementation of Rate4Site^36^ that accounts for recombination breakpoints, using an LG substitution matrix. We calculated the Robinson-Foulds distance between our maximum-likelihood phylogenies for J and its specificity region using *ete3-compare^37^*. We divided the Robinson-Foulds distance (2276.00) by its maximum possible value (2412.00) to get the normalized distance (0.94).

We calculated gap entropy at each position as: − [*p · log*_2_ (*p*) + (1 − *p*) · *log*_2_(1*−p*)], where *p* is the frequency of gap characters at that position, and 1 -*p* is the frequency of all amino acids at that position.

For all non-parametric bootstrap calculations, we used 100,000 bootstraps, and chose groups of sites without replacement. For example, when testing for high entropy in the 17 MAGE residues, we generated an empirical null distribution by randomly choosing 17 different residues for one sample, and resampling 100,000 times. We used the same procedure for our evolutionary-rate tests.

### Data Availability

All data and analysis code for this project are available at the Dryad digital repository [doi: pending publication]

## Acknowledgments

We thank John Ingraham, Adam Riesselman, Kelly Brock, David Ding, and Alita Burmeister for helpful discussions. K.L.P. is supported by the ELSI Origins Network (EON), which is funded by the John Templeton Foundation. The ideas expressed in this publication are those of the authors and not necessarily those of the funding sources.

## Author contributions

R.M. designed and conceived the study. D.T.J. conducted MAGE experiments. K.L.P. analyzed initial MAGE data. R.M. and S.S. performed statistical and phylogenetic analyses. R.M., D.S.M., and J.R.M. wrote the paper and everyone edited it.

---

**Supplementary Table S1.**
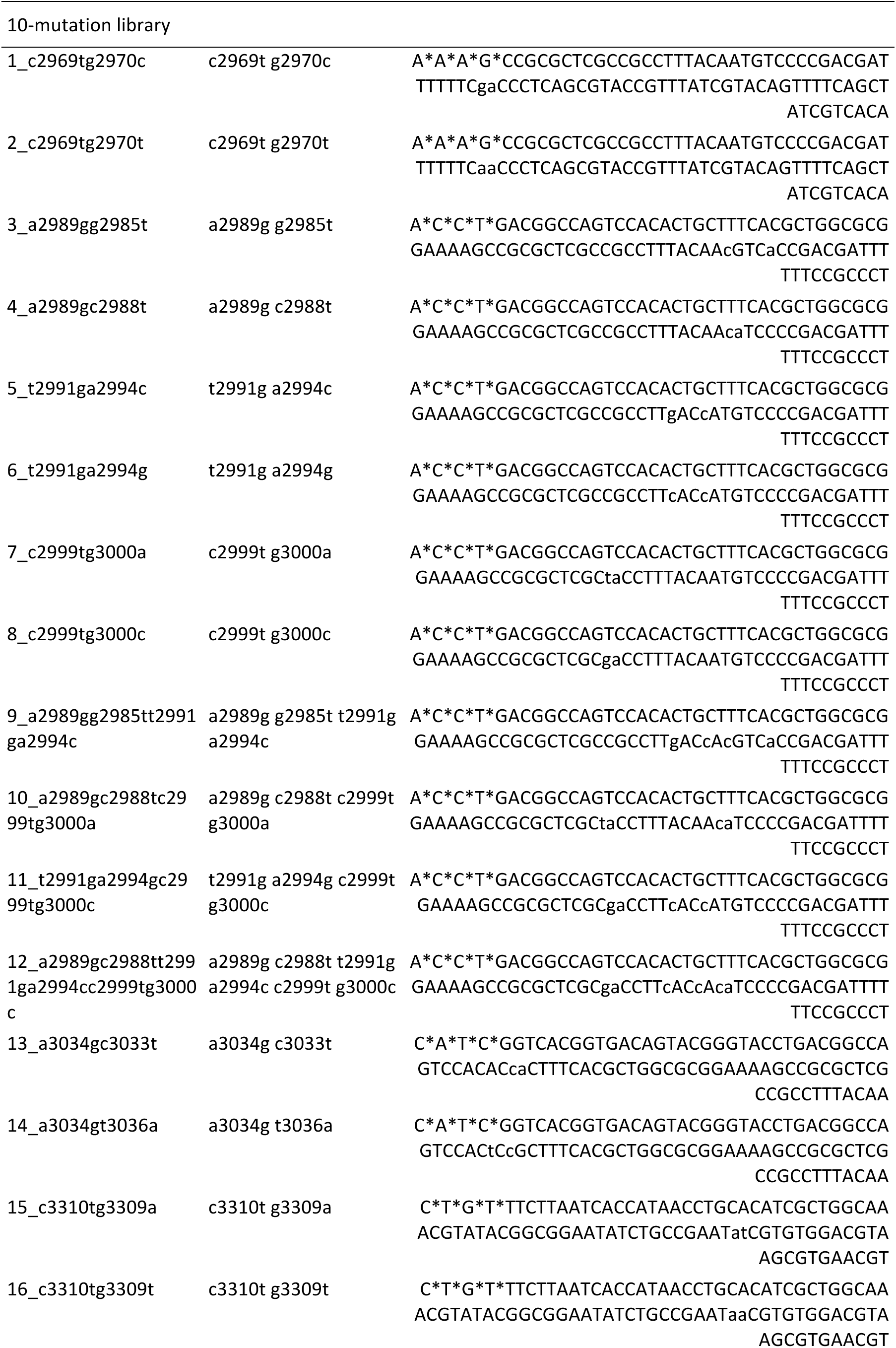
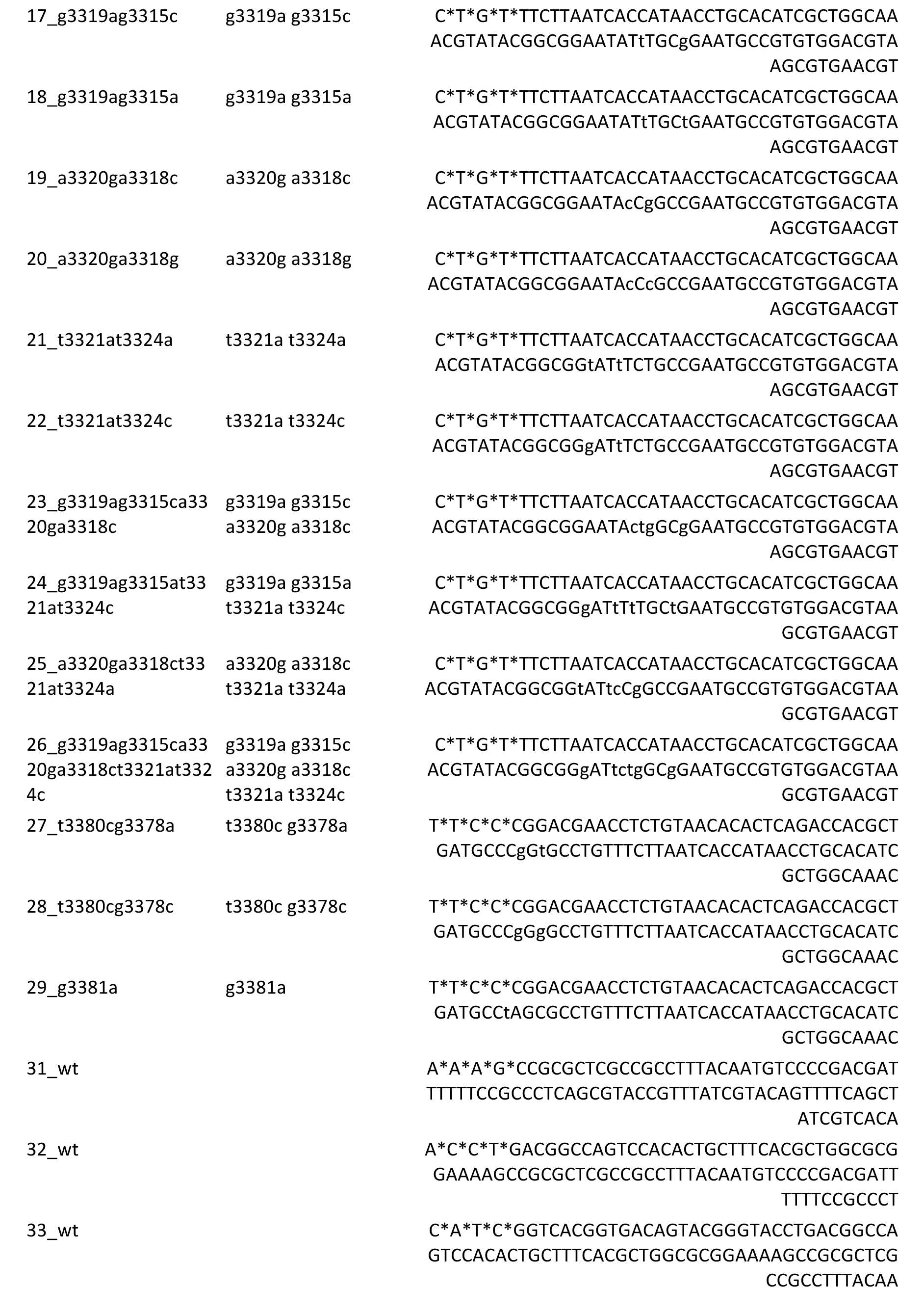
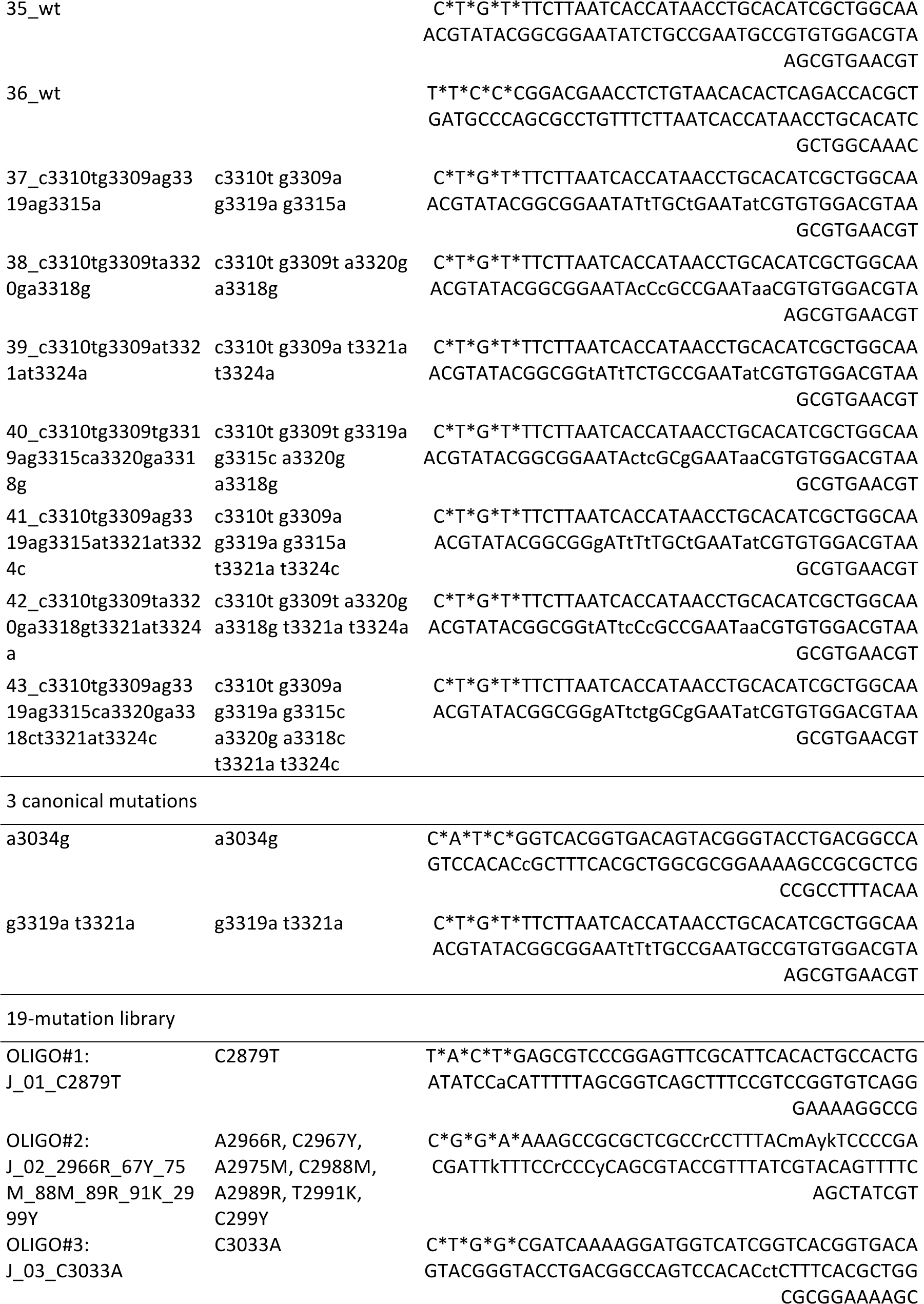
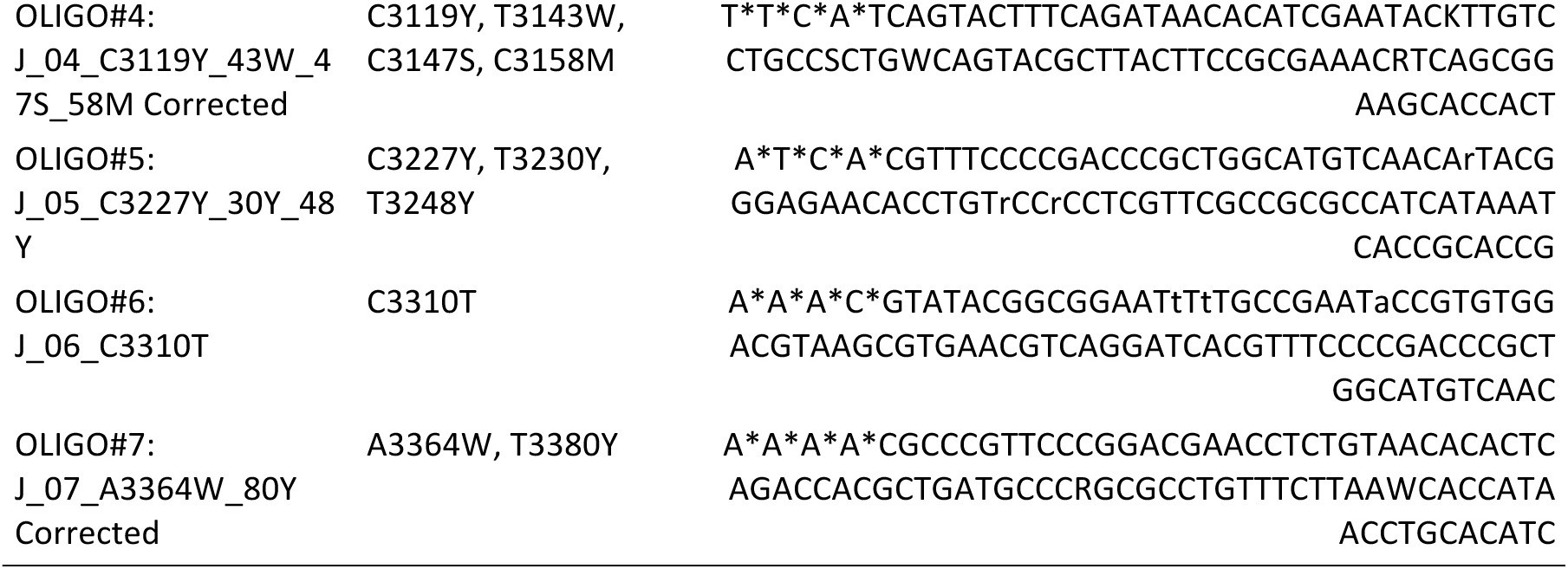
Oligonucleotides (oligos) used to edit λ genomes. Oligos were designed slightly differently for the two libraries. For the first, we inserted a synonymous change near each target. This was done as a silent tag to improve our confidence that an edit had been made successfully. We did not do this for the second library because we found that all intentional nonsynonymous changes were accompanied by the synonymous change. Additionally, for the first library, when mutations fell within 90 bases, we designed a separate oligo for each possible mutation combination. For the second library, we ordered a mixed pool of oligos synthesized that had all combinations of the mutations. Asterisks indicate phosphorothioated bases.

**Supplementary Table S2.** MAGE experiment design for 19-mutation library.

| **Oligos** | **No.**<br>**cycles** | **No.**<br>**replicates** | **No.**<br>**plaques**<br>**sequenced** | **Notes** |
| --- | --- | --- | --- | --- |
| All 7 | 1 | 2 | 16 | This was a pilot study |
| All 7 | 2 | 2 | 15 | This was a pilot study |
| OLIGO#1 | 1 | 1 | 0 | No plaques formed on *lamB*<sup>-</sup> lawn |
| OLIGO#2 | 1 | 1 | 4 |  |
| OLIGO#3 | 1 | 1 | 0 | No plaques formed on *lamB*<sup>-</sup> lawn |
| OLIGO#4 | 1 | 1 | 0 | No plaques formed on *lamB*<sup>-</sup> lawn |
| OLIGO#5 | 1 | 1 | 4 |  |
| OLIGO#6 | 1 | 1 | 0 | No plaques formed on *lamB*<sup>-</sup> lawn |
| OLIGO#7 | 1 | 1 | 0 | No plaques formed on *lamB*<sup>-</sup> lawn |
| All 7 | 1 | 3 | 10 |  |
| All 7 | 2 | 3 | 12 |  |
| All 7 | 3 | 3 | 8 |  |
| All 7 | 4 | 3 | 10 |  |
| All 7 | 5 | 3 | 15 |  |
| All 7 | 6 | 3 | 13 |  |
| All 7 | 7 | 3 | 11 |  |
| All 7 | 8 | 3 | 11 |  |

**Supplementary Figure S1.**
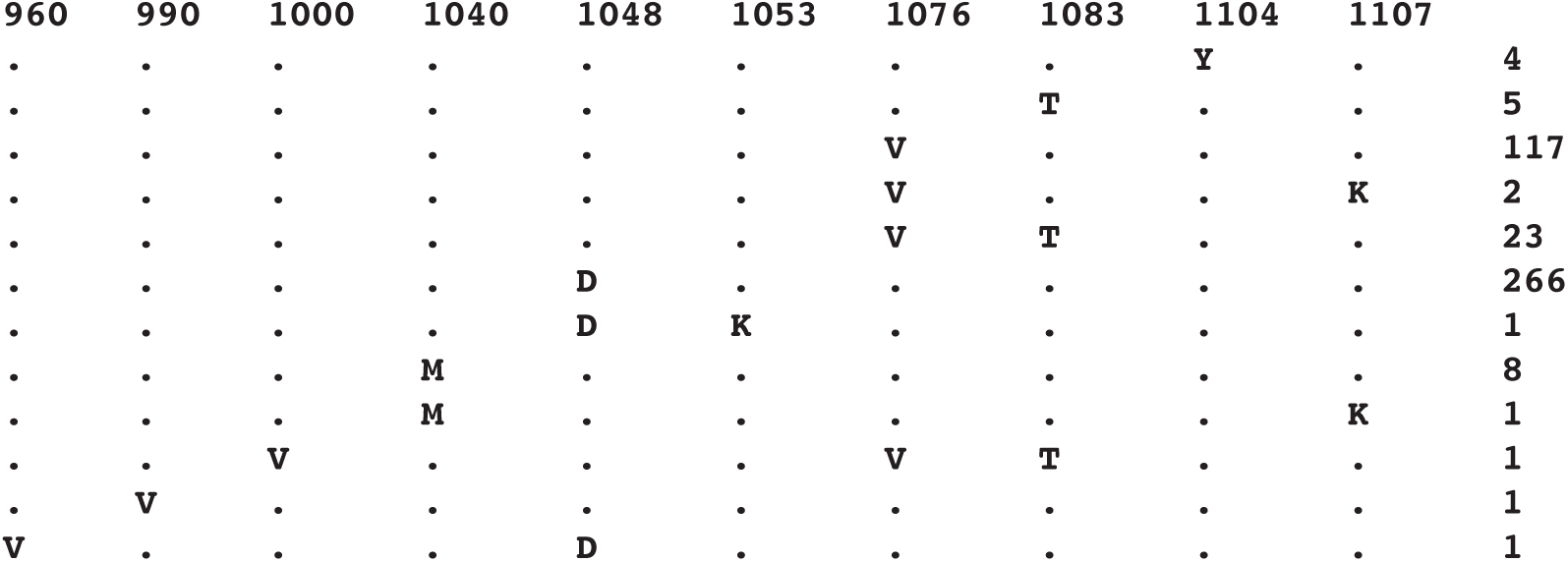
Occurrence of engineered MAGE substitutions in 1207 natural J sequences.

**Supplementary Data File 1 |** Alignment of 1207 full-length J homologs analyzed in this paper.

**Supplementary Data File 2 |** Spreadsheet of gap entropies for each site in the alignment of 1207 full-length J homologs.

